# Joint estimation of contamination, error and demography for nuclear DNA from ancient humans

**DOI:** 10.1101/022285

**Authors:** Fernando Racimo, Gabriel Renaud, Montgomery Slatkin

**Author notes:** Email address (Fernando Racimo). These authors contributed equally to this work (Fernando Racimo, Gabriel Renaud).

## Abstract

When sequencing an ancient DNA sample from a hominin fossil, DNA from present-day humans involved in excavation and extraction will be sequenced along with the endogenous material. This type of contamination is problematic for downstream analyses as it will introduce a bias towards the population of the contaminating individual(s). Quantifying the extent of contamination is a crucial step as it allows researchers to account for possible biases that may arise in downstream genetic analyses. Here, we present an MCMC algorithm to co-estimate the contamination rate, sequencing error rate and demographic parameters – including drift times and admixture rates – for an ancient nuclear genome obtained from human remains, when the putative contaminating DNA comes from present-day humans. We assume we have a large panel representing the putative contaminant population (e.g. European, East Asian or African). The method is implemented in a C++ program called ’Demographic Inference with Contamination and Error’ (DICE). We applied it to simulations and genome data from ancient Neanderthals and modern humans. With reasonable levels of genome sequence coverage (> 3X), we find we can recover accurate estimates of all these parameters, even when the contamination rate is as high as 50%.

## 1. Author Summary

When extracting and sequencing ancient DNA from human remains, a recurrent problem is the presence of DNA from the paleontologists, archaeologists or geneticists that may have handled the fossil. If a DNA library is highly contaminated, this will introduce biases in downstream analyses, so it is important to determine the amount of extraneous DNA. Different methods exist for this purpose, but few are applicable to the nuclear genome, and none of them can extract reliable genomic information from highly contaminated samples. Thus, samples with high rates of contamination are usually discarded. Here, we present a method to jointly estimate contamination and error rates, along with demographic parameters, like drift times and admixture rates. Our method can serve to uncover important details about the evolutionary history of archaic and early modern humans from ancient DNA samples, even if those samples are highly contaminated.

## 2. Introduction

When sequencing a human genome using ancient DNA (aDNA) recovered from fossils, a common practice is to assess the amount of present-day human contamination in a sequencing library [1, 2, 3, 4, 5, 6]. Several methods exist to obtain a contamination estimate. First, one can look at ‘diagnostic positions’ in the mitochondrial genome at which a particular archaic population may be known to differ from all present-day humans. Then, one counts how many aDNA fragments support the present-day human base at those positions. This is the most popular technique and has been routinely deployed in the sequencing of Neanderthal genomes [7, 1]. However, contamination levels of the mitochondrial genome may sometimes differ drastically from those of the nuclear genome [8, 9].

A second technique involves assessing whether the sample was male or female using the number of fragments that map to the X and the Y chromosomes. After determining the biological sex, the proportion of reads that are non-concordant with the sex of the archaic individual are used to estimate contamination from individuals of the opposite sex (e.g. Y-chr reads in an archaic female genome are indicative of male contamination) [8, 1, 10, 4]. Another method uses a maximum-likelihood approach to estimate contamination, but is only applicable to single-copy chromosomes, like the X chromosome in individuals known *a priori* to be male [11, 12]. Finally, one last technique involves using a maximum-likelihood approach to co-estimate the amount of contamination, sequencing error and heterozygosity in the entire autosomal nuclear genome [1, 3], using an optimization algorithm such as L-BFGS-B [13].

Afterwards, if the aDNA library shows low levels of present-day human contamination (< ∼2%), demographic analyses are performed on the sequences while ignoring the contamination. If the library is highly contaminated, it is usually treated as unusable and discarded. Neither of these outcomes is optimal: contaminating fragments may affect downstream analyses, while discarding the library as a whole may waste precious genomic data that could provide important demographic insights.

One way to address this problem was proposed by Skoglund et al. [14], who developed a statistical framework to separate contaminant from endogenous DNA fragments by using the patterns of chemical deamination characteristic of ancient DNA. The method produces a score which reflects the odds that a particular fragment is endogenous or not. This approach, however, may not be able to make a clean distinction between the two sources of DNA, especially for young ancient DNA samples, as chemical degradation may not have affected all fragments belonging to the ancient individual.

Instead of (or in addition to) attempting to separate the two type of fragments before performing a demographic analysis, one could incorporate the uncertainty stemming from the contaminant fragments into a probabilistic inference framework. Such an approach has already been implemented in the analysis of a haploid mtDNA archaic genome [15]. However, mtDNA represents a single gene genealogy, and, so far, no equivalent method has been developed for the analysis of the nuclear genome, which contains the richest amount of population genetic information. Here, we present a method to co-estimate the contamination rate, per-base error rate and a simple demography for an autosomal nuclear genome of an ancient hominin. We assume we have a large panel representing the putative contaminant population, for example, European, Asian or African 1000 Genomes data [16]. The method uses a Bayesian framework to obtain posterior estimates of all parameters of interest, including population-size-scaled divergence times and admixture rates.

## 3. Methods

### 3.1. Basic framework for estimation of error and contamination

We will first describe the probabilistic structure of our inference framework. We begin by defining the following parameters:

- *r_c_:* contamination rate in the ancient DNA sample coming from the contaminant population
- *∊*: error rate, i.e. probability of observing a derived allele when the true allele is ancestral, or vice versa.
- *i*: number of chromosomes that contain the derived allele at a particular site in the ancient individual (*i* = 0, 1 *or* 2)
- *dj*: number of derived fragments observed at site *j*
- **d**: vector of *d_j_* counts for all sites *j* = {1,…, *N*} in a genome
- *a_j_*: number of ancestral fragments observed at site *j*
- **a**: vector of *a_j_* counts for all sites *j* = {1,…, *N*} in a genome
- *W_j_*: known frequency of a derived allele in a candidate contaminant panel at site *j* (0 ≤ *w_j_* ≤ 1)
- **w**: vector of *w_j_* frequencies for all sites *j* = {1,…, *N*} in a genome
- *K*: number of informative SNPs used as input
- *θ*: population-scaled mutation rate. *θ* = 4*N_e_μ*, where *N_e_* is the effective population size and *μ* is the per-generation mutation rate.

We are interested in computing the probability of the data given the contamination rate, the error rate, the derived allele frequencies from the putative contaminant population (**w**) and a set of demographic parameters (**Ω**). We will use only sites that are segregating in the contaminant panel and we will assume that we observe only ancestral or derived alleles at every site (i.e. we ignore triallelic sites). In some of the analyses below, we will also assume that we have additional data (**O**) from present-day populations that may be related to the population to which the sample belongs. The nature of the data in **O** will be explained below, and will vary in each of the different cases we describe. The parameters contained in **Ω** may simply be the population-scaled times separating the contaminant population and the sample from their common ancestral population. However, **Ω** may include additional parameters, such as the admixture rate – if any – between the contaminant and the sample population. The number of parameters we can include in **Ω** will depend on the nature of the data in **O**.

For all models we will describe, the probability of the data can be defined as:

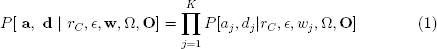

where

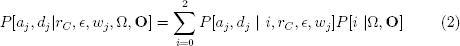

Here, *i* is the true (unknown) genotype of the ancient sample, and *P*[*i* |Ω, **O**] is the probability of genotype *i* given the demographic parameters and the data.

We focus now on computation on the likelihood for one site *j* in the genome. In the following, we abuse notation and drop the subscript *j*. Given the true genotype of the ancient individual, the number of derived and ancestral fragments at a particular site follows a binomial distribution that depends on the genotype, the error rate and the rate of contamination [1, 3]:

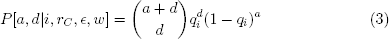

where

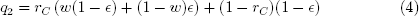

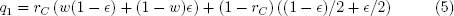

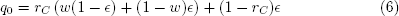

In the sections below, we will turn to the more complicated part of the model, which is obtaining the probability *P*[*i*|**Ω**, **O**] for a genotype in the ancient sample, given particular demographic parameters and additional data available. We will do this in different ways, depending on the kind of data we have at hand.

### 3.2. Diffusion-based likelihood for neutral drift separating two populations

First, we will work with the case in which **O** = **y**, where **y** is a vector of frequencies *y_j_* from an “anchor” population that may be closely related to the population of the ancient DNA sample. An example of this scenario would be the sequencing of a Neanderthal sample that is suspected to have contamination from present-day humans, from which many genomes are available.

For all analyses below, we restrict to sites where 0 < *y_j_* < 1. Note that it is entirely possible (but not required) that **y** = **w**, meaning that, aside from the ancient DNA sample, the only additional data we have are the frequencies of the derived allele in the putative contaminant population, which we can use as the anchor population too. However, it is also possible to use a contaminant panel that is different from the anchor population (Figure 1.A). We will assume we have sequenced a large number of individuals from a panel of the contaminant population (for example, The 1000 Genomes Project panel) and that the panel is large enough such that the sampling variance is approximately 0. In other words, the frequency we observe in the contaminant panel will be assumed to be equal to the population frequency in the entire contaminant population. In this case, **Ω** = {*τ*_C_, *τ*_A_}, where *τ_A_* and *τ_C_* are defined as follows:

**Figure 1.**
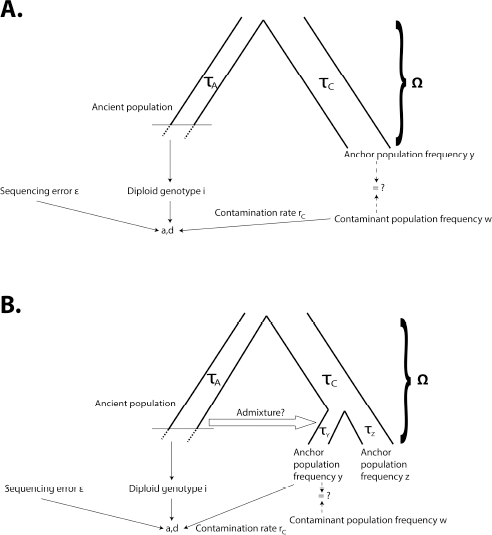
A) Schematic of two-population modeling framework: at each site, derived and ancestral fragments (a, d) are binomially sampled from the true genotype of the archaic individual, with some amount of contamination and error. In turn, the true genotype depends on a demographic model, which can include the contaminant population. B) Schematic of three-population modeling framework, incorporating admixture between the archaic population and one of two anchor populations.

*τ_A_*: drift time (i.e. time in generations scaled by twice the haploid effective population size) separating the population to which the ancient individual belongs from the ancestor of both populations

*τ_C_*: drift time separating the anchor population from the ancestor of both populations

We need to calculate the conditional probabilities *P*[*i*|**Ω**, **O**] = **P**[**i**|**y**, *τ*_C_, *τ*_A_] for all three possibilities for the genotype in the ancient individual: *i* = 0, 1 or 2. To obtain these expressions, we rely on Wright-Fisher diffusion theory (reviewed in Ewens [17]), especially focusing on the two-population site-frequency spectrum (SFS) [18]. The full derivations can be found in Appendix A, and lead to the following formulas:

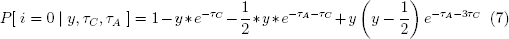

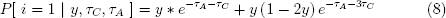

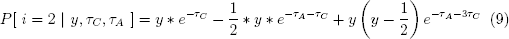

We generated 10,000 neutral simulations using msms [19] for different choices of *τ*_C_ and *τ*_A_ (with *θ* = 20 in each simulation) to verify our analytic expressions were correct (Figure 2). The probability does not depend on *θ*, so the choice of this value is arbitrary.

**Figure 2.**
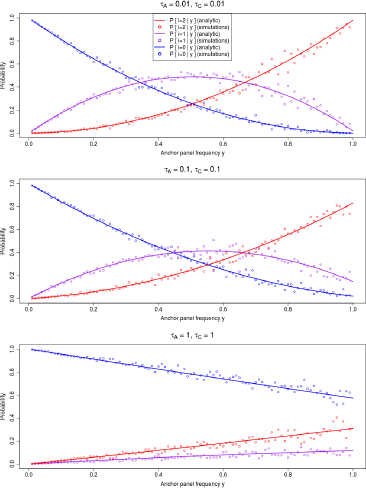
Comparison of analytic solutions to *P*[*i*|*y*, *τ_C_*, *τ_A_*] and simulations under neutrality from msms, for different choices of *τ_A_* and *τ_C_*.

The above probabilities allows us to finally obtain *P*[*i* | *y_j_*, **Ω**, **O**].

### 3.3. Estimating drift and admixture in a three-population model

Although the above method gives accurate results for a simple demographic scenario, it does not incorporate the possibility of admixture from the ancient sample to the contaminant population. This is important, as the signal of contamination may mimic the pattern of recent admixture. We will assume that, in addition to the ancient DNA sample, we also have the following data, which constitute **O**:

1. A large panel from a population suspected to be the contaminant in the ancient DNA sample. The sample frequencies from this panel will be labeled **w**, as before.
2. Two panels of genomes from two “anchor” populations that may be related to the ancient DNA sample. One of these populations – called population Y – may (but need not) be the same population as the contaminant and may (but need not) have received admixture from the ancient population (Figure 1.B). The sample frequencies for this population will be labeled as **y**. The other population – called Z – will have sample frequencies labeled **z**. We will assume the drift times separating these two populations are known (parameters *τ_Y_* and *τ_Z_* in Figure 1.B). This is a reasonable assumption as these parameters can be accurately estimated without the need of using an ancient outgroup sample, as long as admixture is not extremely high.

We can then estimate the remaining drift parameters, the error and contamination rates and the admixture time (*β*) and rate (α) between the archaic population and modern population *Y*. The diffusion solution for this three-population scenario with admixture is very difficult to obtain analytically. Instead, we use a numerical approximation, implemented in the program *∂a∂i* [20].

### 3.4. Markov Chain Monte Carlo method for inference

We incorporated the likelihood functions defined above into a Markov Chain Monte Carlo (MCMC) inference method, to obtain posterior probability distributions for the contamination rate, the sequencing error rate, the drift times and the admixture rate. Our program – which we called ’DICE’ – is coded in C++ and is freely available at: http://grenaud.github.io/dice/. We assumed uniform prior distributions for all parameters, and the boundaries of these distributions can be modified by the user.

For the starting chain at step 0, an initial set of parameters *X*_0_ = {*r*_*C*0_, ∊_0_, Ω_0_} is sampled randomly from their prior distributions. At step *k*, a new set of values for step *k* + 1 is proposed by drawing values for each of the parameters from normal distributions. The mean of each of those distributions is the value for each parameter at state *X_k_* and the standard deviation is the difference between the upper and lower boundary of the prior, divided by a constant that can be increased or decreased to achieve a desired rate of acceptance of new states [21]. By default, this constant is equal to 1, 000 for all parameters. The new state is accepted with probability:

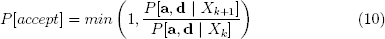

where *P*[**a**, **d** | *X_k_*] is the likelihood defined in Equation 1.

Unless otherwise stated below, we ran the MCMC chain for 100,000 steps in all analyses, with a burn-in period of 40,000 and sampling every 100 steps. The sampled values were then used to construct posterior distributions for each parameter.

### 3.5. Multiple error rates and ancestral state misidentification

Fu et al. [5] showed that, when estimating contamination, ancient DNA data can be better fit by a two-error model than a single-error model. In that study, the authors co-estimate the two genome-wide error rates along with the proportion of the data that is affected by each rate. Therefore, we also included this error model as an option that the user can choose to incorporate when running our program.

Furthermore, we developed an alternative error estimation method that allows the user to flag transition polymorphisms, which are more likely to have occurred due to cytosine deamination in ancient DNA. These sites are therefore likely to be subject to different error rates than those common in present-day sequencing data [22, 23]. Our program can then estimate two error rates separately: one for transitions and one for transversions. Finally, we incorporated an option to include an ancestral state misidentification (ASM) parameter, which should serve to correct for mispolarization of alleles [24].

### 3.6. BAM file functionality

The standard input for DICE is a file containing counts of particular ancestral/derived base combinations and SNP frequencies (see README file online). As an additional feature, we also developed a module for the user to directly input a BAM file and a file containing population allele frequencies for the anchor and contaminant panels, rather than the standard input. The user can either choose to convert the BAM file to native DICE format using a program provided with the software package and then run the program, or run it directly on the BAM file. In the latter case, instead of calculating genome-wide error parameters, the program will calculate error parameters specific to each sequenced fragment, based on mapping qualities, base qualities and estimated deamination rates at each site (see Appendix B).

## 4. Results: two-population method

### 4.1. Simulations

We first used DICE to obtain posterior distributions from simulated data, under the two-population inference framework. We simulated two populations (i.e. an archaic and a modern human population) with constant population size that split a number of generations ago. For each demographic scenario tested, we generated 20,000 independent replicates (theta=1) in *ms* [25], making sure each simulation had at least one usable SNP. In general, this yielded ∼80,000 usable SNPs in total. We then proceeded to sample derived and ancestral allele counts using the same binomial sampling model we use in our inference framework, under different sequencing coverage and contamination conditions. In all simulations, the contaminant panel was the same as the anchor population panel. We then applied our method to the combined set of ∼80,000 SNPs.

Figure 3 and 4 show parameter estimation results from various demographic and contamination scenarios for a low-coverage (3X) and a high-coverage (30X) archaic genome, respectively, with low sequencing error (0.1%), and a contaminant/anchor population panel of 100 haploid genomes. In both cases, the method accurately estimates the error rate, the contamination rate and the drift parameters. All parameters are also accurately estimated for the same scenarios even if the sequencing error rate is high (10%) (Figure S1).

**Figure 3.**
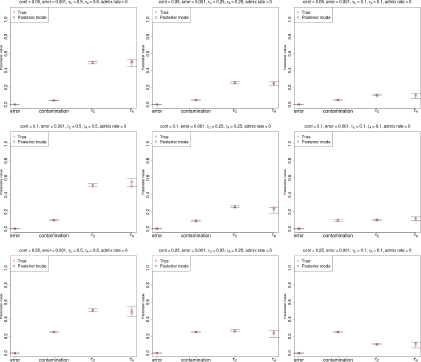
Estimation of parameters for a low-coverage ancient DNA genome (3X) with low sequencing error (0.1%), no admixture and a large anchor population panel (100 haploid genomes). Error bars represent 95% posterior intervals.

**Figure 4.**
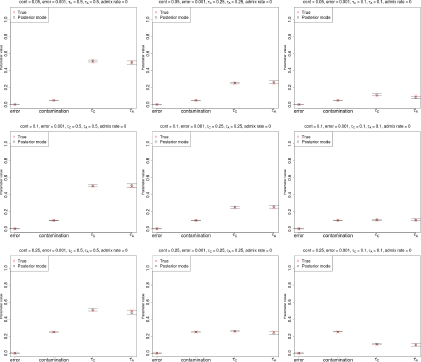
Estimation of parameters for a high-coverage ancient DNA genome (30X) with low sequencing error (0.1%), no admixture and a large anchor population panel (100 haploid genomes). Error bars represent 95% posterior intervals.

Figures 5, S2, S3, S4 show how well the method does at estimating parameters over a wide range of contamination and drift scenarios, by displaying the absolute difference between simulated parameters and their corresponding posterior modes. So long as coverage is high (for example, 5X or 30X), the contamination and anchor drift parameters are accurately estimated even at 75% contamination. The method performs well even if the drift times on both sides of the tree are as small as ≈ 0.001 or as large as ≈ 5, but starts becoming inaccurate when contamination is extremely high. In general, the contamination rate and anchor drifts are easier to determine than the drift corresponding to the ancient population.

**Figure 5.**
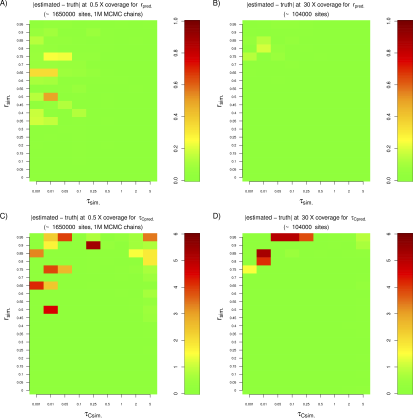
We tested the performance of the two-population method under a variety of drift and contamination scenarios for a sample of very low (0.5X) or very high (30X) coverage. We found that we needed more sites (≈ 1:6 million) to obtain accurate estimates from the low coverage sample. The MCMC chain was also run for a longer time (1 million steps). The top row shows the absolute difference between the estimated and the simulated contamination rate, while the bottom row shows the absolute difference corresponding to the anchor drift. In all simulations, the anchor drift was set to be equal to the ancient sample drift.

We find that for samples of very low coverage (0.5X, 1X, 1.5X) we require a larger number of sites to obtain accurate estimates (Figures S5, S6, S7). For example, for a sample of 0.5X coverage, we tried different numbers of independent replicate simulations and found that at 800,000 replicates, we obtained approximately 1.6 million valid SNPs for inference, which was enough to reach reasonable levels of accuracy (Figure S14). We note that this number of SNPs is approximately the same as what is available, for example, in the low-coverage (0.5X) Mezmaiskaya Neanderthal genome [4], which contains about 1.55 million valid sites with coverage ≥ 1, and which we analyze below. We also observed that the MCMC chain in some of these simulations needed a longer time to converge than when testing samples of higher coverage, especially when contamination is very high, and so in this set of simulations, we ran it for 1 million steps instead of 100,000, with a burn-in of 940,000 steps and sampling every 100 steps. Finally, we note that our failure to recover the true parameters under low coverage in a single MCMC run is partly due to the chain failing to converge. Indeed, when we run the MCMC 10 times and recover the estimates from the chain with the highest posterior probability, we are able to obtain increased accuracy relative to the single run, especially when the drift parameters are extremely low and when the contamination rate is extremely high (Figures S8, S9, S10).

Finally, we tested the method on simulations in a more realistic scenario, in which we generated ancient and contaminant fragments based on empirical fragment sizes and then mapped them to a simulated reference genome using BWA [26] with default parameters. We produced DNA sequences from the output of msms [19] via seq-gen v.1.3.3 [27] with the HKY substitution model [28]. This allows for multiple substitutions to occur at the same site since the split from chimpanzee (which could cause ASM). We then simulated ancient DNA fragments that had a fragment size distribution emulating empirical distributions. Contaminant fragments were also sampled from the contaminant population. We used the deamination rates from the singlestranded library from the Loschbour ancient individual [29] (∼ 8% at the 5’ end and ∼ 34% at the 3’ end with a residual deamination rate of ∼ 1% along the whole fragment) to artificially deaminate the ancient fragments. We simulated sequencing errors on both the ancient and contaminant fragments using empirical sequencing error rates from a PhiX library (Illumina Corp.) sequenced at the Max Planck Institute for Evolutionary Anthropology on an Illumina HiSeq, basecalled using freeIbis [30]. With the same empirical PhiX dataset distribution, we generated quality scores for each nucleotide. Fragments were mapped back to a random individual from the contaminant panel. Figure 6 shows DICE’s performance on this scenario with different error models. In all cases, we find that the parameters are estimated with high accuracy. As expected, the ts/tv model infers a higher error rate at transitions, due to the additional errors introduced by deamination on the ends of the ancient fragments.

**Figure 6.**
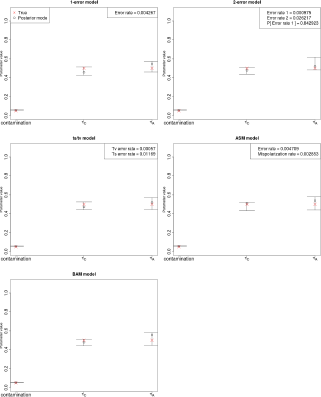
Estimation of parameters for a high-coverage ancient DNA genome (30X) simulated under a realistic scenario in which fragments from the ancient and contaminant genome were generated and then mapped to a reference genome. We allowed for multiple substitutions at the same site after the split from chimp, as well as sequencing errors and post-mortem deamination errors at the ends of the fragments. The five panels show results from inferring parameters under five different error rate models. Top-left: single-error model. Top-right: two-error model [5]. Middle-left: model with separate errors for transitions (ts) and tranversions (tv). Middle-right: single-error model with an ancestral state misidentification parameter. Bottom-left: Model in which errors were inferred individually at each site, using base and mapping qualities obtained from the simulated BAM file. Error bars represent 95% posterior intervals.

### 4.2. Performance under violations of model assumptions

We evaluated the consequences of different violations of model assumptions. We started by observing the effects of using a small modern human panel. Figure S12 shows results for cases in which the contaminant/anchor panel is made up of only 20 haploid genomes. In this case, all parameters are estimated accurately, with only a slight bias towards overestimating the drift parameters, presumably because the low sampling of individuals acts as a population bottleneck, artificially increasing the drift time parameters estimated.

Additionally, we simulated a scenario in which only a single human contaminated the sample. That is, rather than drawing contaminant fragments from a panel of individuals, we randomly picked a set of two chromosomes at each unlinked site and only drew contaminant fragments from those two chromosomes. Figure S13 shows that inference is robust to this scenario, unless the contamination rate is very high (25%). In that case, the drift of the archaic genome is substantially under-estimated, but the error, contamination and anchor drift parameters only show slight inaccuracies in the estimate.

We then investigated the effect of admixture in the anchor/contaminant population from the archaic population, occurring after their divergence, which we did not account for in the simple, two-population model (Figure S11). In this case, the error and the contamination rates are accurately estimated, but both drift times are underestimated. This is to be expected, as admixture will tend to homogenize allele frequencies and thereby reduce the apparent drift separating the two populations.

### 4.3. Identifying the contaminant population

We sought to see whether we would use our method to identify the contaminant population, from among a set of candidate contaminants (for example, different present-day human panels). Because our MCMC samples are samples from the posterior distribution of the parameters and not the marginal likelihood of the data over the entire parameter space, we cannot perform proper Bayesian model selection. Instead, we used the posterior mode as a heuristic statistic that may suggest which panel is most likely to have contaminated the sample. We validated this choice of statistic using simulations under a variety of demographic scenarios (Figure S15). We simulated 5-population trees of varying drift times. The outgroup was chosen to be the ancient population and the rest were chosen to be the present-day human populations (A, B, C and D). One of the populations (A) was the true contaminant. To add another layer of complexity, we also allowed for admixture (at 0%, 5% and 50% rate) from the ancient population to the ancestral population of A and B. We then ran our MCMC method four times on each of these demographic scenarios, using D as the anchor and different panels as the putative contaminant in each run.

Figure S16 shows that the lowest posterior mode always corresponds to the run that uses the true contaminant (A), and that the mode decreases the farther the tested contaminant is from the true contaminant in the tree. Additionally, Figures S17, S18, S19 show the effect of misspecifying the contaminant panel for different admixture scenarios. The error rate and the anchor drift time are correctly estimated, even when the candidate contaminant is highly diverged from the true contaminant, while the other two parameters are more sensitive to misspecification. In general, the correct candidate contaminant produces the highest posterior probability and yields the best parameter estimates.

### 4.4. Empirical data

We first applied our method to published ancient DNA data from a high-coverage genome (52X) from Denisova cave in Siberia (the Altai Neanderthal) [4], and visually ensured that the chain had converged. The demographic, error and contamination estimates are shown in Table 1. We used the African (AFR) 1000 Genomes Phase 3 panel [16] as the anchor population. The drift times estimated for both samples are consistent with the known demographic history of Neanderthals and modern humans, and the contamination rates largely agree with previous estimates (see Discussion below).

**Table 1.**
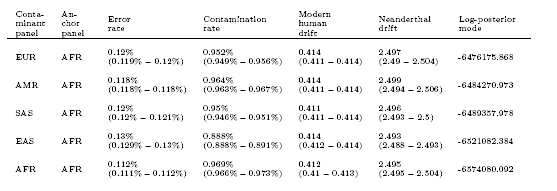
Posterior modes of parameter estimates under the two-population inference framework for the Altai Neanderthal autosomal genome. We used different 1000G populations as candidate contaminants. Africans were the anchor population in all cases, so the modern human drift is with respect to Africans. Values in parentheses are 95% posterior quantiles.

We ran our method with different putative contaminant panels: Africans (AFR), East Asians (EAS), Native Americans (AMR), Europeans (EUR), South Asians (SAS). For the Altai sample, we observe a contamination rate of ∼ 1% and an error rate of ∼ 0.1%, regardless of which panel we use. Furthermore, the drift on the Neanderthal side of the tree seems to be 6 times as large as the drift on the modern human side of the tree, reflecting the smaller effective population size of Neanderthals after their divergence. The EUR panel is the one with the highest posterior mode (Table 1).

We then tested a variety of ancient DNA nuclear genome sequences at different levels of coverage, obtained via different methods (shotgun sequencing and SNP capture) and from different hominin groups (modern humans and Neanderthals). We used AFR as the anchor panel and either AFR (Table S1) or EUR (Table S2) as the contaminant panel. For samples of high and medium average coverage, the MCMC converges to reasonable values for all parameters. For example, we estimate the ancient population drift parameter (*τ_A_*) to be larger in Neanderthals than in various modern humans sampled across Eurasia, as the effective population size of the former was smaller and their split time to Africans was larger.

However, for samples of very low coverage, we observe a failure of some of the parameters to properly converge, as the MCMC seems to get stuck in the boundaries of parameter space. We tested different boundaries and the problem remains. This appears to be less of a problem when using AFR as the putative contaminant panel than when using EUR as the putative contaminant panel, presumably because of the larger amount of SNPs that may be informative for inference. In the former case, we only observe this problem when samples are at lower than ∼ 0.5X coverage. In the latter case, we observe the problem for samples at lower than ∼ 3X coverage.

For example, the low-coverage Neanderthal genome (0.5X) from Mezmaiskaya Cave in Western Russia [4] seems to converge to parameters within the prior boundaries when using AFR as the contaminant panel but the ancient population drift gets stuck in the upper limit of parameter space when any of the other panels are used as contaminants (Table S3). Regardless of which contaminant panel is used, there is good agreement with the modern human drift parameter obtained when using the Altai Neanderthal genome. However, we note that when using non-African populations as the contaminants, we obtain a higher (∼ 5%) contamination rate in the Mezmaiskaya Neanderthal than in the Altai Neanderthal. It is currently unclear to us whether this is due to the MCMC failing to properly converge or to a real feature of the data.

We sought to determine the robustness of our results to different levels of GC content. We did this because we initially hypothesized that endogenous DNA might be preserved at lower rates when GC content is low, leading to the presence of proportionally more contaminant DNA. We partitioned the Altai Neanderthal genome into three different regions of low (0% – 30%), medium (31% – 69%) and high (70% – 100%) GC content, using the ’GC content’ track downloaded from the UCSC genome browser [31]. We then used the two-population method to infer contamination, error and drift parameters, using Africans as the anchor population and Europeans as the contaminant population (Figure S20). We observe that contamination rates are higher in low-GC regions than in medium-GC regions (Welch one-sided t-test on the posterior samples, P < 2.2e-16), which in turn have higher contamination rates than high-GC regions (P < 2.2e-16). The opposite trend occurs in the error estimates, while the drift parameters are largely unaffected. However, we find that the differences we observe across GC levels are almost entirely eliminated by removing CpG sites from the input dataset (Figure S20), as CpG sites are known to have higher mutation rates than the rest of the genome. For this reason, we recommend filtering them out when testing for contamination on ancient DNA datasets, which is what was done in Tables 1 and 2.

**Table 2.** Posterior modes of parameter estimates under the three-population inference framework for the Altai Neanderthal autosomal genome. We used different 1000G populations as candidate contaminants. In all cases, Africans were the unadmixed anchor population and Europeans were the admixed anchor population. The ancestral human drift refers to the drift in the modern human branch before the split of Europeans and Africans. The post-split European-specific and African-specific drifts were estimated separately without the archaic genome (*τ_Afr_* = 0.009, *τ_Eur_* = 0.255).

| Contaminant panel | Unadmixed anchor panel | Admixed anchor panel | Error rate | Contamination rate | Ancestral human drift | Neanderthal drift | Admixture rate | Log-posterior mode |
| --- | --- | --- | --- | --- | --- | --- | --- | --- |
| EUR | AFR | EUR | 0.119%<br>(0.119% – 0.12%) | 0.967%<br>(0.954% – 0.967%) | 0.411<br>(0.405 – 0.414) | 2.669<br>(2.656 – 2.689) | 1.72%<br>(1.682% – 1.805%) | -7452958.125 |
| AMR | AFR | EUR | 0.119%<br>(0.118% – 0.12%) | 0.967%<br>(0.962% – 0.974%) | 0.407<br>(0.402 – 0.412) | 2.677<br>(2.651 – 2.708) | 1.661%<br>(1.618% – 1.696%) | -7461041.325 |
| SAS | AFR | EUR | 0.122%<br>(0.122% – 0.123%) | 0.95%<br>(0.944% – 0.955%) | 0.399<br>(0.398 – 0.406) | 2.682<br>(2.677 – 2.695) | 1.469%<br>(1.422% – 1.48%) | -7465214.726 |
| EAS | AFR | EUR | 0.13%<br>(0.129% – 0.132%) | 0.896%<br>(0.884% – 0.903%) | 0.421<br>(0.413 – 0.428) | 2.702<br>(2.658 – 2.706) | 2.388%<br>(2.009% – 2.447%) | -7509504.053 |
| AFR | AFR | EUR | 0.117%<br>(0.117% – 0.119%) | 0.957%<br>(0.945% – 0.964%) | 0.409<br>(0.409 – 0.418) | 2.681<br>(2.66 – 2.702) | 1.837%<br>(1.766% – 1.961%) | -7554080.773 |

Finally, we tested a present-day Yoruba genome (HGDP00936) sequenced to high coverage [4], which should not contain any contamination. Indeed, when applying our method, we find this to be the case (Figure S21). We infer 0% contamination, regardless of whether we use EUR or AFR as the candidate contaminant. Furthermore, the anchor drift time is very close to 0 when using AFR as the anchor population (as the sample belongs to that same population), while it is non-zero (= 0.22) when using EUR, which is consistent with the drift time separating Europeans from the ancestor of Europeans and their closest African sister populations [32].

## 5. Results: three-population method

### 5.1. Simulations

We applied our three-population method to estimate both drift times and admixture rates. We simulated a high-coverage (30X) archaic human genome under various demographic and contamination scenarios. Each of the two anchor population panels contained 20 haploid genomes. The admixture time was 0.08 drift units ago, which under a constant population size of 2N=20,000 would be equivalent to 1,600 generations ago. When running our inference program, we set the admixture time prior boundaries to be between 0.06 and 0.1 drift units ago.

We find that the admixture time is inaccurately estimated under this implementation – likely due to lack of information in the site-frequency spectrum – so we do not show estimates for that parameter below. For admixture rates of 0%, 5% or 20%, the error and contamination parameters are estimated accurately in all cases (Figures S22, S23 and S24, respectively). The method is less accurate when estimating the demographic parameters, especially the admixture rate which is sometimes under-estimated. Importantly though, the accuracy of the contamination rate estimates are not affected by incorrect estimation of the demographic parameters.

We also tested what would happen if the admixture time was simulated to be recent: 0.005 drift units ago, or 100 generations ago under a constant population size of 2N=20,000. When estimating parameters, we set the prior for the admixture time to be between 0 and 0.01 drift units ago. In this last case, we observe that the drift times and the admixture rate (20%) are more accurately estimated than when the admixture event is ancient (Figure 7).

**Figure 7.**
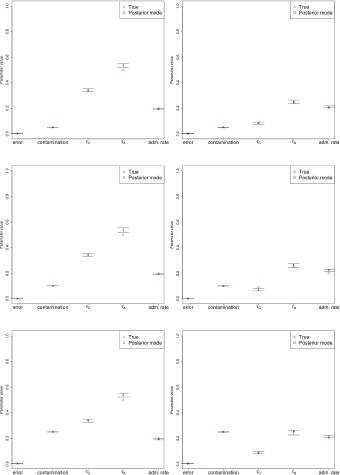
Estimation of error, contamination and demographic parameters in various three-population demographic scenarios, where the admixture rate is 20% and the admixture time was recent (0.005 drift units ago). The prior used for the admixture time was uniform over [0,0.01]. Error bars represent 95% posterior intervals.

As before, we also verified that the posterior mode was a good proxy to identify the true contaminant (A), when running the MCMC using different contaminant panels (A, B, C and D). In all cases, we used D as the unadmixed anchor panel and B as the admixed anchor panel. Results are shown in Figure S25 for all the demographic scenarios from Figure S15. Again, we observe that the true contaminant (A) is always the one that corresponds to the lowest posterior probability, though we again caution that because we do not have the marginal probabilities, we cannot formally perform model selection to favor a particular panel. Furthermore,the admixture rate from the ancient population into the ancestors of A and B is robustly estimated unless the true contaminant (A) is highly diverged from the candidate contaminant (Figures S26, S27, S28, for admixture rates of 0%, 5% and 50%, respectively).

### 5.2. Empirical data

We also applied the three-population inference framework to the high-coverage Altai Neanderthal genome. We first estimated the two drift times specific to Europeans and Africans after the split from each other (*τ_Y_* and *τ_Z_*, respectively), using ∂a∂i and the L-BFGS-B likelihood optimization algorithm [13], but without using the archaic genome (*τ_Afr_* = 0.009, *τ_Eur_* = 0.255). Then, we used our MCMC method to estimate the rest of the drift times, the archaic admixture rate and the contamination and error parameters in the Neanderthal genome. We set the admixture time prior boundaries to be between 0.06 and 0.1 drift units ago, which is a realistic time frame given knowledge about modern human – Neanderthal cohabitation in Eurasia [33]. The error rate and contamination rates we obtain are similar to those obtained under the two-population method, and we estimate an admixture rate from Neanderthals into modern humans of 1.72% for the choice of contaminant panel with the highest posterior mode – which is again EUR (Table 2).

We also applied the method to the low-coverage Mezmaiskaya Neanderthal genome. As before, we are able to reach convergence for all parameters (including the admixture rate) with the exception of the Neanderthal drift, which gets stuck in the upper boundary of parameter space (Table S4).

## 6. Discussion

We have developed a new method to jointly infer demographic parameters, along with contamination and error rates, when analyzing an ancient DNA sample. The method can be deployed using a C++ program (DICE) that is easy to use and freely downloadable. We therefore expect it to be highly applicable in the field of paleogenomics, allowing researchers to derive useful information from previously unusable (highly contaminated) samples, including archaic humans like Neanderthals, as well as ancient modern humans.

Applications to simulations show that the error and contamination parameters are estimated with high accuracy, and that demographic parameters can also be estimated accurately so long as enough information (e.g. a large panel of modern humans) is available. The drift time estimates reflect how much genetic drift has acted to differentiate the archaic and modern populations since the split from their common ancestral population, and can be converted to divergence times in generations if an accurate history of population size changes is also available (for example, via methods like PSMC, [34]). Although we cannot perform proper model testing, we found via extensive simulations that the posterior mode of an MCMC run was a robust heuristic statistic to help detect which panel was most likely to have contaminated the sample. We caution, however, that the fact that a particular panel yields a higher posterior mode than another is no guarantee that it is a better fit to the data for demographic scenarios that may be different from the ones we simulated.

We also applied our method to empirical data, specifically to two Neanderthal genomes at high and low coverage, a present-day high-coverage Yoruba genome, and several ancient genome sequences of varying degrees of coverage, some obtained via shotgun-sequencing and some via SNP capture. For the high-coverage Yoruba genome, we infer no contamination, as would be expected from a modern-day sample, and drift times indicating the Yoruba sample indeed belongs to an African population.

The contamination and sequencing error estimates we obtained for the Altai Neanderthal are roughly in accordance with previous estimates [4]. The drift times we obtain under the three-population model for the African population (*τ_C_* + *τ_Afr_*) are approximately 0.411 + 0.009 = 0.42 drift units. The geometric mean of the history of population sizes from the PSMC results in Prüfer et al. [4] give roughly that *N_e_* ≈ 21, 818 since the African population size history started differing from that of Neanderthals, assuming a mutation rate of 1.25 * 10^−8^ per bp per generation. If we assume a generation time of 29 years, and use our drift time in the equation relating divergence time in generations to drift time (*t*/(2*N_e_*) ≈ *τ*), this gives an approximate human-Neanderthal population divergence time of 531,486 years. This number roughly agrees with the most recent estimates obtained via other methods [4]. Additionally, the Neanderthal-specific drift time is approximately 6.5 times as large as the modern human drift time, which is expected as Neanderthals had much smaller population sizes than modern humans [35, 4]. The admixture rate from archaic to modern humans that we estimate is 1.72%, which is consistent with the rate estimate obtained via methods that do not jointly model contamination (1.5 – 2.1%) [4]. In the case of the Altai Neanderthal, we observe that the sample was probably contaminated by one or more individuals with European ancestry.

When testing modern human and Neanderthal ancient genomes of lower coverage than the Altai Neanderthal, we obtain reasonable parameter estimates for samples of medium to high-coverage. However, we run into problems in estimation when the samples are of low coverage. For these reasons, and from our simulation results, we recommend that our method should be used on nuclear genomes with > 3X coverage. The method may converge under certain conditions at coverages as low as 0.5X (for example, in the case of the Mezmaiskaya genome under the two-population model when using AFR as the anchor and contaminant panel), but, in such cases, we caution the user to check convergence is achieved before drawing any conclusions from the estimates. For SNP capture data, we obtain reliable estimates for samples with a minimum coverage of 500,000 sites that are polymorphic in the anchor panel.

The demographic models used in our approach are simple, involving no more than three populations and a single admixture event. This is partly due to limitations of known theory about the diffusion-based likelihood of an arbitrarily complex demography for the 2-D site-frequency spectrum – in the case of the two-population method – and to the inability of ∂a∂i [20] to handle more than 3 populations at a time. In recent years, several studies have made advances in the development of methods to compute the likelihood of an SFS for larger numbers of populations using coalescent theory [36, 37, 38], with multiple population size changes and admixture events. We hope that some of these techniques could be incorporated in future versions of our inference framework.

## 7. Acknowledgments

We thank Kelley Harris, Philip Johnson, Graham Coop, Nicolas Duforet-Frebourg, Joshua Schraiber, Sergi Castellano, Christoph Theunert, Janet Kelso, Rasmus Nielsen and members of the Slatkin and Nielsen labs for helpful advice and discussions.

## Supporting Information

**Figure S1.**
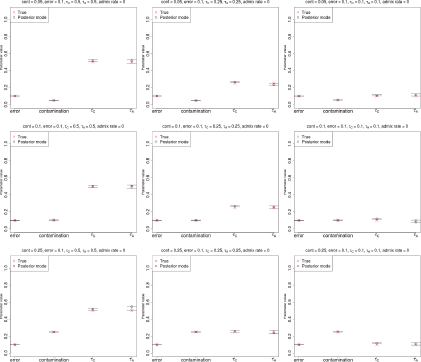
Estimation of parameters for a high-coverage ancient DNA genome (30X) with high sequencing error (10%), no admixture and a large anchor population panel (100 haploid genomes). Error bars represent 95% posterior intervals.

**Table S1.** We applied the two-population method to ancient Neanderthal and modern human genomes ranging from 52X to 0.054X coverage. We tested both shotgun-sequencing data and SNP capture data. We used AFR as both the anchor panel and the putative contaminant panel. Samples are sorted by decreasing mean coverage. We define Convergence to be true (T) if all the parameters stably converged in a region of parameter space that does not include the upper parameter boundary. Otherwise Convergence is false (F). A line separates the two Convergence classes. SNPs = number of SNPs overlapping with anchor panel. Observations = total number of base observations analyzed. SC = SNP capture. SS = shotgun sequencing. HG = hunter-gatherer. LBK = Linear Pottery culture. MN = Middle Neolithic. LN = Late Neolithic. NEA = Neanderthal. MH = Modern Human. LogPos = Log-posterior mode. Reported Cov. = Mean read coverage reported in corresponding study. For SNP capture, this is the mean coverage of the targeted SNPs.

| ID | Study | Group | Type | Description | Reported Cov. | SNPs | Observations | Convergence | Error | Cont. (AFR) | $\tau_G$ (AFR) | $\tau_A$ | LogPos |
| --- | --- | --- | --- | --- | --- | --- | --- | --- | --- | --- | --- | --- | --- |
| Altai | [4] | NEA | SS | Altai Nea. | 52 | 9500771 | 495741350 | T | 0.11% | 0.97% | 0.412 | 2.495 | -6574080.092 |
| Loschbour | [29] | MH | SS | Loschbour | 22 | 8733958 | 181642481 | T | 0.17% | 0.00% | 0.025 | 0.634 | -5905688.289 |
| Stuttgart | [39] | MH | SS | LBK | 19 | 8720170 | 157538109 | T | 0.14% | 0.00% | 0.019 | 0.392 | -5921770.389 |
| I0100 | [39] | MH | SC | LBK | 6.727 | 1017124 | 4608980 | T | 0.09% | 0.00% | 0.025 | 0.361 | -483569.9795 |
| I0061 | [39] | MH | SC | Karelia (HG) | 5.272 | 729066 | 3189601 | T | 0.11% | 0.00% | 0.026 | 0.438 | -394527.9859 |
| I0104 | [39] | MH | SC | Corded Ware (LN) | 4.184 | 912245 | 2714837 | T | 0.13% | 0.00% | 0.022 | 0.325 | -423668.0231 |
| I0406 | [39] | MH | SC | Spain (MN) | 3.947 | 545379 | 3204204 | T | 0.12% | 0.00% | 0.024 | 0.367 | -341002.1352 |
| I0014 | [39] | MH | SC | Motala (HG) | 2.709 | 497524 | 2164912 | T | 0.12% | 0.05% | 0.031 | 0.445 | -295108.9014 |
| Kennewick | [40] | MH | SS | Kennewick | 1.6 | 5725599 | 9648018 | T | 0.90% | 2.13% | 0.021 | 0.603 | -1951145.65 |
| MA-1 | [41] | MH | SS | Mal'ta | 1 | 6969896 | 13578653 | T | 0.26% | 3.64% | 0.026 | 0.603 | -2375835.916 |
| I0111 | [39] | MH | SC | Bell Beaker (LN) | 0.731 | 300636 | 456149 | T | 0.20% | 0.16% | 0.033 | 0.386 | -127455.6442 |
| I0013 | [39] | MH | SC | Motala (HG) | 0.657 | 349019 | 788739 | T | 0.24% | 2.20% | 0.033 | 0.464 | -179713.5404 |
| Mezmaiskaya | [4] | NEA | SS | Mezmaiskaya Nea. | 0.48 | 4896677 | 6811727 | T | 0.52% | 0.01% | 0.406 | 1.756 | -889165.6704 |
| I0439 | [39] | MH | SC | Yamnaya | 0.26 | 176088 | 194152 | F | 0.28% | 15.60% | 0.04 | 3.495 | -61178.22162 |
| I0060 | [39] | MH | SC | Bell Beaker (LN) | 0.105 | 67741 | 73195 | F | 0.24% | 12.03% | 0.045 | 3.344 | -24561.12309 |
| I0804 | [39] | MH | SC | Unetice | 0.054 | 34069 | 35522 | F | 0.32% | 7.23% | 0.042 | 1.566 | -11980.98331 |

**Table S2.**
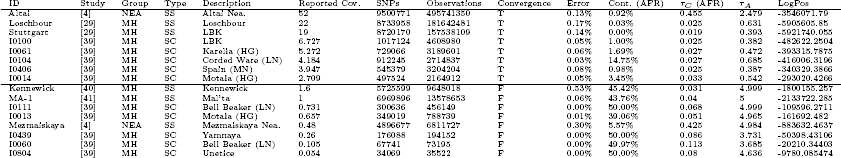
We applied the two-population method to ancient Neanderthal and modern human genomes ranging from 52X to 0.054X coverage. We tested both shotgun-sequencing data and SNP capture data. We used AFR as the anchor panel and EUR as the putative contaminant panel. Samples are sorted by decreasing mean coverage. We define Convergence to be true (T) if all the parameters stably converged in a region of parameter space that does not include the upper parameter boundary. Otherwise Convergence is false (F). A line separates the two Convergence classes. SNPs = number of SNPs overlapping with anchor panel. Observations = total number of base observations analyzed. SC = SNP capture. SS = shotgun sequencing. HG = hunter-gatherer. LBK = Linear Pottery culture. MN = Middle Neolithic. LN = Late Neolithic. NEA = Neanderthal. MH = Modern Human. LogPos = Log-posterior mode. Reported Cov. = Mean read coverage reported in corresponding study. For SNP capture, this is the mean coverage of the targeted SNPs.

**Table S3.**
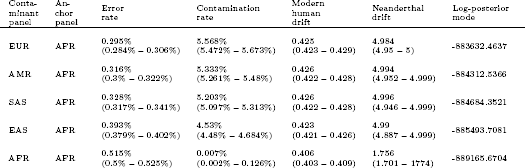
Posterior modes of parameter estimates under the two-population inference framework for the Mezmaiskaya Neanderthal autosomal genome. We used different 1000G populations as candidate contaminants. AFR were the anchor population in all cases, so the modern human drift is with respect to Africans. Values in parentheses are 95% posterior quantiles. Except when using AFR as the contaminant, the Neanderthal drift parameter gets stuck at the upper boundary (5 drift units) of parameter space.

**Table S4.** Posterior modes of parameter estimates under the three-population inference framework for the Mezmaiskaya Neanderthal autosomal genome. We used different 1000G populations as candidate contaminants. In all cases, Africans were the unadmixed anchor population and Europeans were the admixed anchor population. The ancestral human drift refers to the drift in the modern human branch before the split of Europeans and Africans. The post-split European-specific and African-specific drifts were estimated separately without the archaic genome (*τ_Afr_* = 0.009, *τ_Eur_* = 0.255). In all cases, the Neanderthal drift parameter gets stuck at the upper boundary (5 drift units) of parameter space.

| Contaminant panel | Unadmixed anchor panel | Admixed anchor panel | Error rate | Contamination rate | Ancestral human drift | Neanderthal drift | Admixture rate | Log-posterior mode |
| --- | --- | --- | --- | --- | --- | --- | --- | --- |
| AFR | AFR | EUR | 0.517%<br>(0.502% – 0.526%) | 4.663%<br>(4.564% – 4.787%) | 0.428<br>(0.426 – 0.432) | 4.999<br>(4.989 – 5) | 1.609%<br>(1.585% – 1.63%) | -1025944.516 |
| EAS | AFR | EUR | 0.71%<br>(0.697% – 0.721%) | 2.471%<br>(2.403% – 2.564%) | 0.415<br>(0.412 – 0.418) | 4.997<br>(4.985 – 5) | 1.486%<br>(1.462% – 1.508%) | -1028456.347 |
| AMR | AFR | EUR | 0.727%<br>(0.71% – 0.733%) | 2.288%<br>(2.208% – 2.361%) | 0.414<br>(0.412 – 0.417) | 4.999<br>(4.985 – 5) | 1.482%<br>(1.459% – 1.501%) | -1028866.312 |
| SAS | AFR | EUR | 0.724%<br>(0.709% – 0.732%) | 2.315%<br>(2.219% – 2.375%) | 0.414<br>(0.412 – 0.418) | 4.998<br>(4.984 – 5) | 1.479%<br>(1.458% – 1.5%) | -1028823.568 |
| EUR | AFR | EUR | 0.761%<br>(0.745% – 0.77%) | 1.875%<br>(1.784% – 1.928%) | 0.413<br>(0.41 – 0.415) | 4.998<br>(4.984 – 2.5) | 1.463%<br>(1.457% – 1.495%) | -1029429.156 |

**Figure S2.**
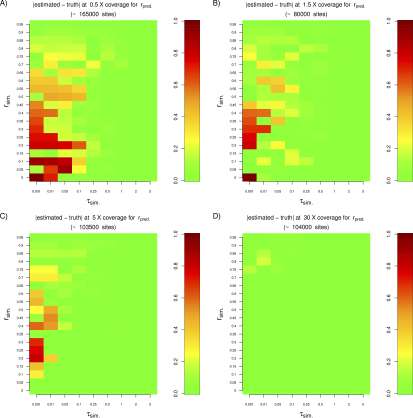
Absolute difference between estimated and simulated contamination rates for a variety of anchor drift and contamination scenarios, for different levels of coverage. In all simulations, the anchor drift was set to be equal to the ancient sample drift.

**Figure S3.**
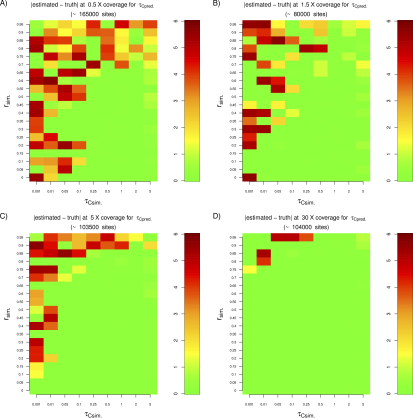
Absolute difference between estimated and simulated anchor drifts for a variety of anchor drift and contamination scenarios, for different levels of coverage. In all simulations, the anchor drift was set to be equal to the ancient sample drift.

**Figure S4.**
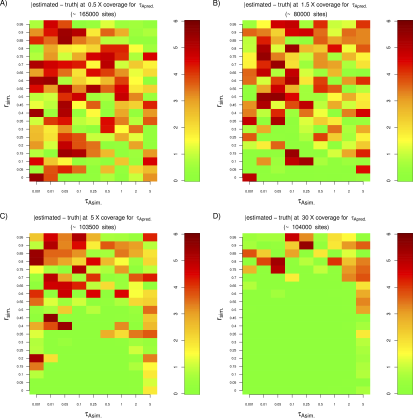
Absolute difference between estimated and simulated ancient sample drifts for a variety of anchor drift and contamination scenarios, for different levels of coverage. In all simulations, the anchor drift was set to be equal to the ancient sample drift.

**Figure S5.**
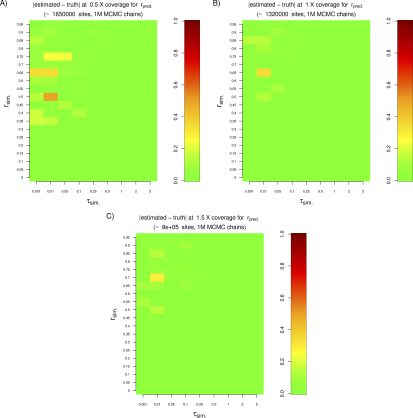
Absolute difference between estimated and simulated contamination rates for a variety of anchor drift and contamination scenarios, when coverage is low (0.5X, 1X or 1.5X). Here, we used a large number of sites and run the MCMC chain for 1 million steps. In all simulations, the anchor drift was set to be equal to the ancient sample drift.

**Figure S6.**
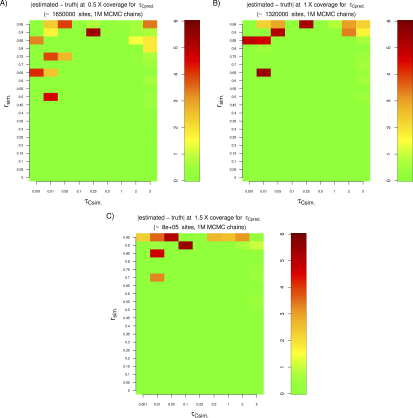
Absolute difference between estimated and simulated anchor drifts for a variety of anchor drift and contamination scenarios, when coverage is low (0.5X, 1X or 1.5X). Here, we used a large number of sites and run the MCMC chain for 1 million steps. In all simulations, the anchor drift was set to be equal to the ancient sample drift.

**Figure S7.**
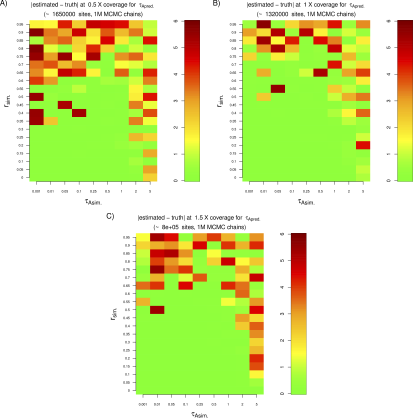
Absolute difference between estimated and simulated ancient sample drifts for a variety of anchor drift and contamination scenarios, when coverage is low (0.5X, 1X or 1.5X). Here, we used a large number of sites and run the MCMC chain for 1 million steps. In all simulations, the anchor drift was set to be equal to the ancient sample drift.

**Figure S8.**
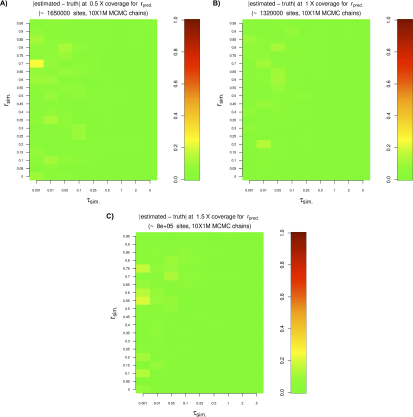
Absolute difference between estimated and simulated contamination rates for a variety of anchor drift and contamination scenarios, when coverage is low (0.5X, 1X or 1.5X). We used a large number of sites and run 10 MCMC chains for 1 million steps each. To ensure convergence, we then selected the chain with the highest posterior probability, and here show estimates from that chain. In all simulations, the anchor drift was set to be equal to the ancient sample drift.

**Figure S9.**
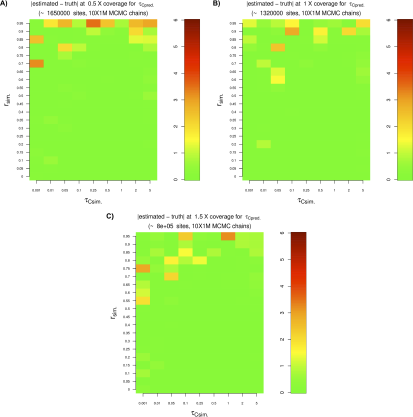
Absolute difference between estimated and simulated anchor drifts for a variety of anchor drift and contamination scenarios, when coverage is low (0.5X, 1X or 1.5X). We used a large number of sites and run 10 MCMC chains for 1 million steps each. To ensure convergence, we then selected the chain with the highest posterior probability, and here show estimates from that chain. In all simulations, the anchor drift was set to be equal to the ancient sample drift.

**Figure S10.**
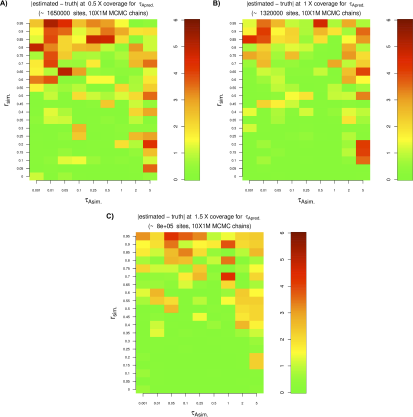
Absolute difference between estimated and simulated ancient sample drifts for a variety of anchor drift and contamination scenarios, when coverage is low (0.5X, 1X or 1.5X). We used a large number of sites and run 10 MCMC chains for 1 million steps each. To ensure convergence, we then selected the chain with the highest posterior probability, and here show estimates from that chain. In all simulations, the anchor drift was set to be equal to the ancient sample drift.

**Figure S11.**
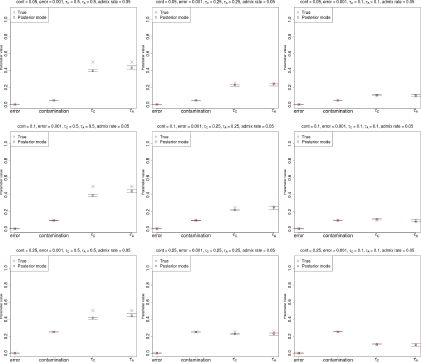
Estimation of parameters for a high-coverage ancient DNA genome (30X) with low sequencing error (0.1%), a large anchor population panel (100 haploid genomes) and admixture in the anchor population from the archaic population (5%), using the two-population inference framework, which does not model admixture. Error bars represent 95% posterior intervals.

**Figure S12.**
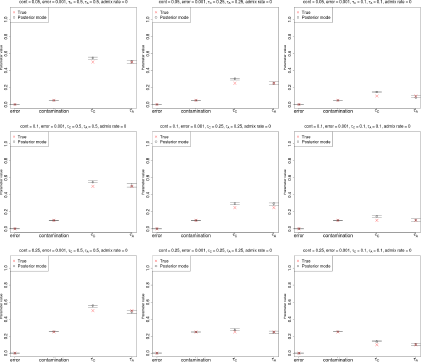
Estimation of parameters for a high-coverage ancient DNA genome (30X) with low sequencing error (0.1%), no admixture and a small anchor population panel (20 haploid genomes). Error bars represent 95% posterior intervals.

**Figure S13.**
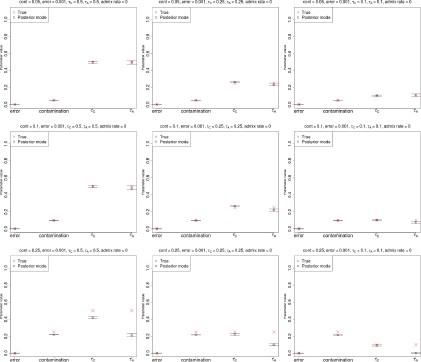
Estimation of parameters for a high-coverage ancient DNA genome (30X), when the contaminant fragments are exclusively drawn from a single diploid individual from the contaminant panel. Error bars represent 95% posterior intervals.

**Figure S14.**
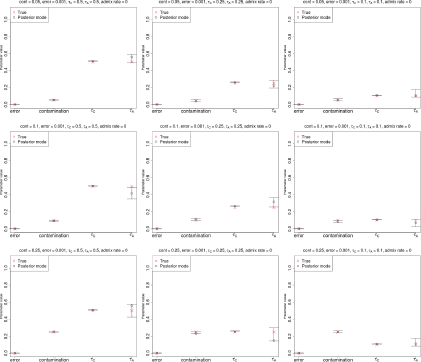
Estimation of parameters for an ancient DNA genome of very low coverage (0.5X) with low sequencing error (0.1%) and a large anchor population panel (100 haploid genomes). Note that unlike the rest of the simulations, the number of SNPs used in this case was approximately 1.6 million instead of 80,000, and the MCMC chain was run for 1 million steps instead of 100,000. Using a lower number of SNPs or running the chain for a shorter time resulted in inaccurate inferences. Error bars represent 95% posterior intervals.

**Figure S15.**
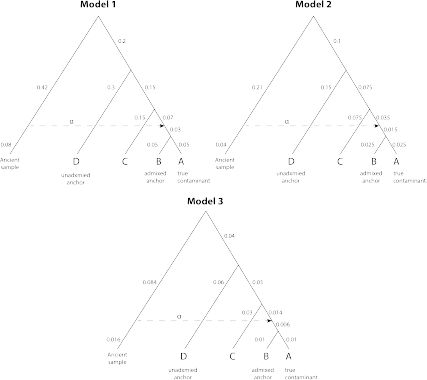
Three demographic models used to test the method when the contaminant is misspecified. When testing the two-population method, we set panel A as the true contaminant and panel D as the anchor. When testing the three-population method, we set panel A as the true contaminant, panel D as the unadmixed anchor and panel B as the admixed anchor. The numbers on the branches represent the drift parameters. The parameter *α* represents the admixture rate from the ancient population into the ancestor of A and B.

**Figure S16.**
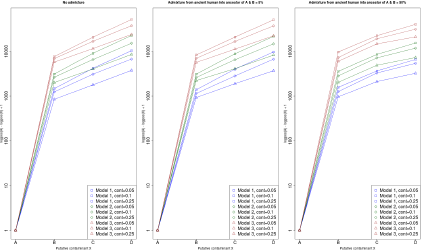
When testing different putative contaminants, the highest mode of the posterior likelihoods from the MCMC under the two-population model corresponds to the true contaminant (panel A). The y-axis shows the difference between the log-posterior for contaminant panel A and the log-posterior for different candidate contaminant panels (A, B, C, D). We added a 1 to the difference to be able to plot the difference on a logarithmic scale. The three panels contain results for three admixture scenarios (from left to right: admixture rate of 0%, 5% and 50%) and each panel shows the difference under different contamination rates and demographic models (see Figure S15).

**Figure S17.**
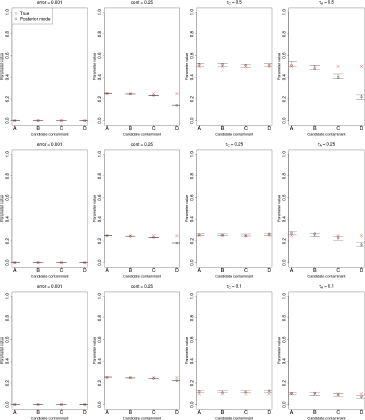
Parameters estimates under the two-population model using different putative contaminants, when the true contaminant is panel A. Each row of panels represents a different set of drift parameters, keeping the contamination rate fixed at 25% and the error rate at 0.1%. In this case, the admixture rate from the ancient population to the ancestor of A and B was kept at 0%. The anchor panel used was panel D (see Figure S15).

**Figure S18.**
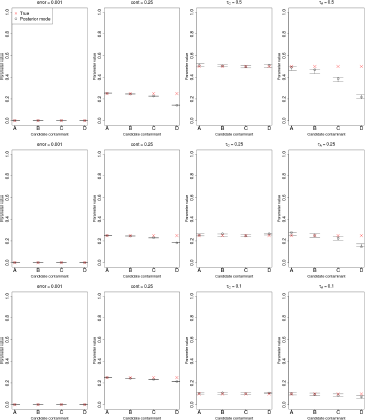
Parameters estimates under the two-population model using different putative contaminants, when the true contaminant is panel A. Each row of panels represents a different set of drift parameters, keeping the contamination rate fixed at 25% and the error rate at 0.1%. In this case, the admixture rate from the ancient population to the ancestor of A and B was kept at 5%. The anchor panel used was panel D (see Figure S15).

**Figure S19.**
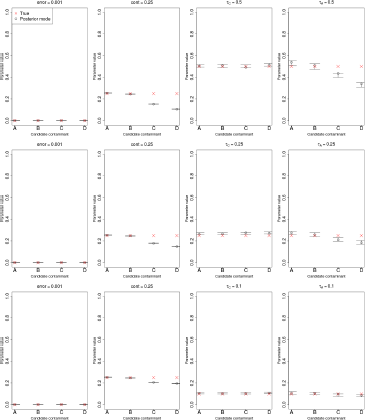
Parameters estimates under the two-population model using different putative contaminants, when the true contaminant is panel A. Each row of panels represents a different set of drift parameters, keeping the contamination rate fixed at 25% and the error rate at 0.1%. In this case, the admixture rate from the ancient population to the ancestor of A and B was kept at 50%. The anchor panel used was panel D (see Figure S15).

**Figure S20.**
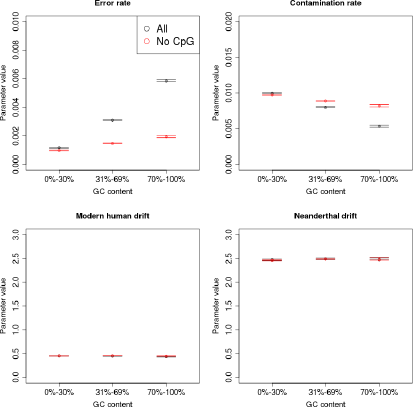
Estimation of parameters for the Altai Neanderthal genome across different GC levels using the two-population model, while keeping (black) or removing (red) CpG sites from the input dataset. Error bars represent 95% posterior intervals.

**Figure S21.**
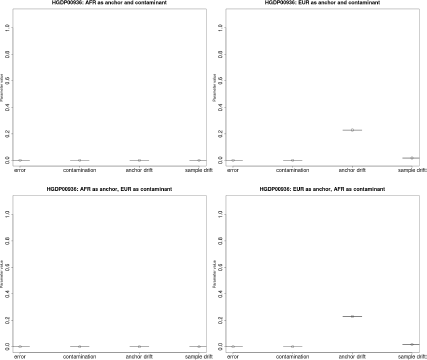
We tested one of the Yoruba genomes from Prüfer et al. [4] and obtain an estimate of 0% contamination, regardless of whether we use Europeans or Africans as the candidate contaminant. The anchor drift time is close to 0 when using Africans as the anchor population, as the sample belongs to that same population, while it is non-zero (= 0.22) when using Europeans. Error bars represent 95% posterior intervals.

**Figure S22.**
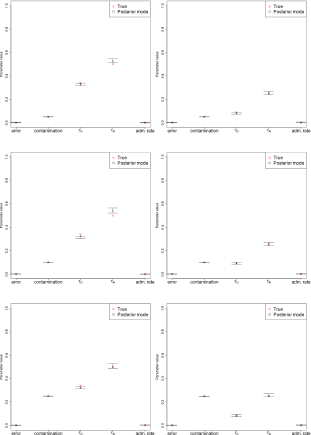
Estimation of error, contamination and demographic parameters in various three-population demographic scenarios, where the admixture rate is 0%. The prior used for the admixture time was uniform over [0.06,0.1]. Error bars represent 95% posterior intervals.

**Figure S23.**
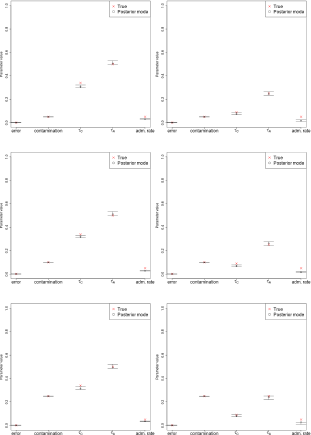
Estimation of error, contamination and demographic parameters in various three-population demographic scenarios, where the admixture rate is 5% and the admixture time is ancient (0.08 drift units ago). The prior used for the admixture time was uniform over [0.06,0.1]. Error bars represent 95% posterior intervals.

**Figure S24.**
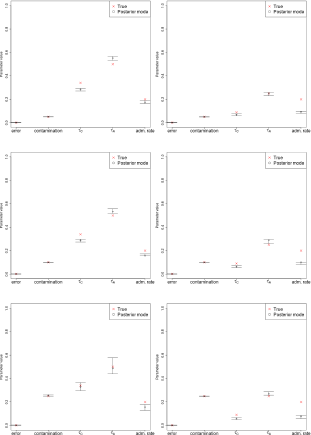
Estimation of error, contamination and demographic parameters in various three-population demographic scenarios, where the admixture rate is 20% and the admixture time is ancient (0.08 drift units ago). The prior used for the admixture time was uniform over [0.06,0.1]. Error bars represent 95% posterior intervals.

**Figure S25.**
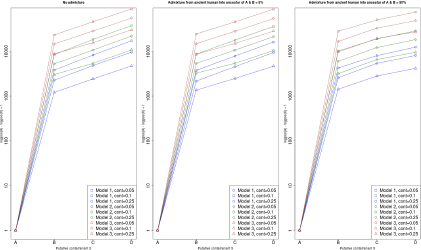
When testing different putative contaminants, the highest mode of the posterior likelihoods from the MCMC under the three-population model corresponds to the true contaminant (panel A). The y-axis shows the difference between the log-posterior for contaminant panel A and the log-posterior for different candidate contaminant panels (A, B, C, D). We added a 1 to the difference to be able to plot the difference on a logarithmic scale. The three panels contain results for three admixture scenarios (from left to right: admixture rate of 0%, 5% and 50%) and each panel shows the difference under different contamination rates and demographic models (see Figure S15).

**Figure S26.**
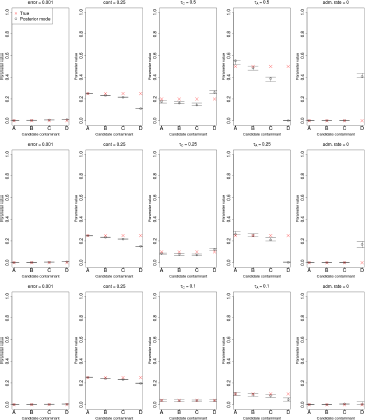
Parameters estimates under the three-population model using different putative contaminants, when the true contaminant is panel A. Each row of panels represents a different set of drift parameters, keeping the contamination rate fixed at 25% and the error rate at 0.1%. In this case, the admixture rate from the ancient population to the ancestor of A and B was kept at 0%. The unadmixed anchor panel used was panel D and the admixed anchor panel was panel B (see Figure S15).

**Figure S27.**
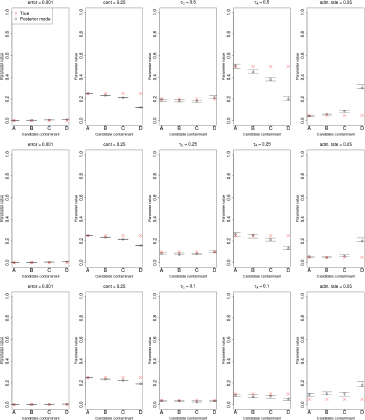
Parameters estimates under the three-population model using different putative contaminants, when the true contaminant is panel A. Each row of panels represents a different set of drift parameters, keeping the contamination rate fixed at 25% and the error rate at 0.1%. In this case, the admixture rate from the ancient population to the ancestor of A and B was kept at 5%. The unadmixed anchor panel used was panel D and the admixed anchor panel was panel B (see Figure S15).

**Figure S28.**
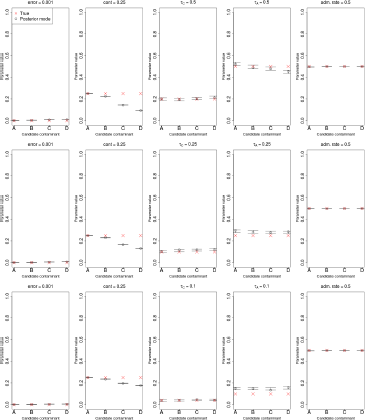
Parameters estimates under the three-population model using different putative contaminants, when the true contaminant is panel A. Each row of panels represents a different set of drift parameters, keeping the contamination rate fixed at 25% and the error rate at 0.1%. In this case, the admixture rate from the ancient population to the ancestor of A and B was kept at 50%. The unadmixed anchor panel used was panel D and the admixed anchor panel was panel B (see Figure S15).

## Appendix A. Genotype probabilities conditional on a demography

Below we derive formulas 7, 8 and 9. Recall that we are interested in calculating the conditional probabilities *P*[*i*|**Ω**, **O**] = **P**[**i**|**y**, *τ*_C_, *τ*_A_] for all three possibilities for the genotype in the ancient individual: *i* = 0, 1 or 2. These can be obtained from the definition of conditional probability. Let 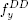 be the joint probability that a site has frequency *y* (0 < *y* < 1) in the contaminant panel and is homozygous for the derived allele in the ancient individual. Let 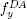 be the joint probability that a site has frequency *y* in the contaminant panel and is heterozygous in the ancient individual. Finally, let 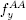 be the joint probability that a site has frequency *y* in the anchor panel and is homozygous for the ancient allele in the ancient individual. Then:

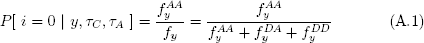

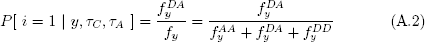

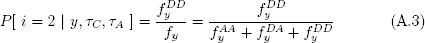

In the above expressions, the functions *f* depend on *T_C_* and *T_A_*, but we omit this conditioning for ease of notation. As can be seen, all we need to find is the joint probabilities 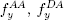 and 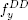. Here is where diffusion theory comes into play. Let *ϕ*(*y, τ*|*x*, 0) be the Kimura solution to the neutral forward diffusion equation in the absence of mutation [42], given a frequency *x* at time 0 and an elapsed drift time *τ*:

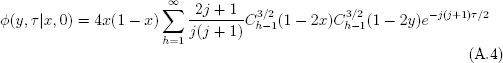

Here, *x* is the unknown population frequency of the derived allele in the ancestral population and 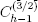 (•) is the Gegenbauer polynomial of order h-1 [43].

Assuming the ancestral population follows an equilibrium frequency distribution *g(x)* = *θ/x*, we can write 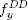 as follows:

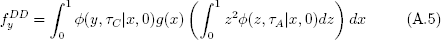

where *z* is the unknown population frequency of a derived allele in the population to which the ancient individual belongs.

The expression in parentheses is the second moment of the transition density and its solution is known [44]:

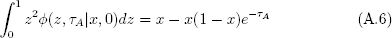

This results in:

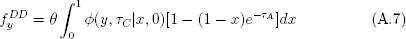

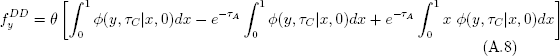

The integral of the first two terms of the sum was solved in Chen et al.

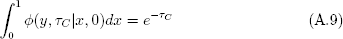

The third term of the sum can be solved by noting that, though the integrand is an infinite sum (i.e. formula A.4 multiplied by *x*), only the integrals of the first two terms of that infinite sum are not equal to 0. This can be seen by integrating the parts of the terms of that infinite sum that depend on *x*:

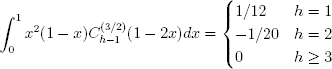

Therefore, after integrating the first two terms of the infinite sum, we obtain:

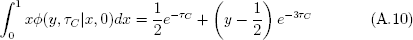

So we finally arrive at:

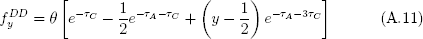

We can obtain 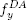 in a similar fashion:

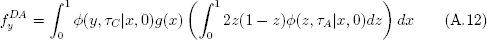

Solving the term in the parentheses:

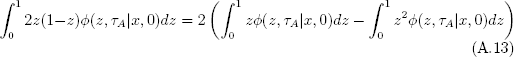

The first term of the difference is the first moment of the transition density, which is equal to *x* [44], while the second term is the second moment (formula A.6). Therefore:

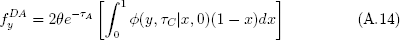

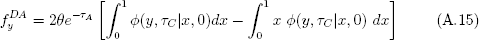

And after using formulas A.9 and A.10, we obtain:

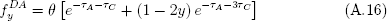

To obtain 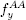, we know that, assuming the anchor population to be at equilibrium:

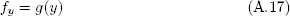

And therefore:

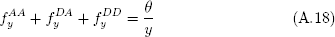

So we finally obtain:

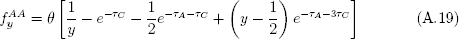

We now have all the elements necessary to obtain the conditional probabilities from formulas A.1, A.2 and A.3, which immediately lead us to formulas 7, 8 and 9.

## Appendix B. Probabilistic inference using BAM files

Here, we briefly explain the way we infer fragment-specific error parameters in the optional BAM mode of DICE. Let ℝ be the set of all fragments in the BAM file, and *R_j_* ∈ ℝ be a particular aligned fragment of length *l*. For fragment *R_j_*, let {*b*_*j*,1_,…, *b_j,l_*} be the individuals nucleotides in the fragment. At each position of the fragment, there is a specific probability *κ_j,i_* that the base is erroneous. This probability is provided by the basecaller. Below, we will compute the likelihood of observing a base *b_j,i_* ∈ *R_j_* under a bi-allelic model, given an error rate *κ_j, i_*. Below, we focus on an individual fragment *R_j_* and an individual position *i* on that fragment, so for simplicity, we drop the subscripts *i* and *j* and we let *b_j, i_* = *b* and *κ_j, i_* = *κ*.

Let *ν* be the base that was originally sampled at a given site, before deamination or mismapping. This base could be ancestral or derived. Let *P_dam_*[*ν* → *b*] be the probability of substitution from v to b due to postmortem chemical damage. The probabilities of different types of damage (e.g. C → or G → A) occurring at different positions of a fragment can be computed following Ginolhac et al. [45] and Jónsson et al. [46], producing a matrix that can be provided to DICE as input. We offer the possibility of specifying different post-mortem damage matrices for the endogenous and the contaminant fragments.

Let *E* denote the event that a sequencing error has occurred, let *D* the event that chemical damage has occurred, let *M* be the event that *R_j_* was correctly mapped and let ¬ denote the complement of an event (i.e. event has not occurred). We define the probability of observing sequenced base *b* given that no sequencing error has occurred at a position on a correctly mapped fragment that was originally *υ*, by summing over two possibilities, either chemical damage occurred or it did not:

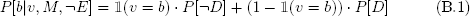

Here, 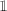(*υ* = *b*) is an indicator function that is equal to 1 if *υ* is equal to b, and 0 otherwise. The probabilities *P[D]* and *P[¬D]* are respectively equal to *P_dam_*[*υ* → *b*] and 1 – *P_dam_*[*υ* → *b*].

Subsequently, we compute *P*[*b*|*υ*, *M*], the probability of observing *b* given *υ* under the assumption that *R_j_* was mapped at the correct genomic location. We have:

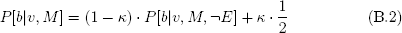

This is because if a sequencing error has occurred, the probability of observing *b* is independent of *υ*, and therefore 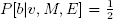. Finally, let *P[M]* be the probability that the fragment *R_j_* is mapped at the correct location as given by the mapping quality. The probability of seeing *b* given that *υ* was the base that was sampled before deamination is then:

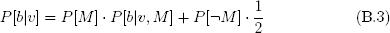

The probability of observing *b* given that the fragment was mismapped is independent of *υ*, hence 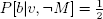. If either the base quality or mapping quality indicate a probability of error of 100%, *P*[*b*|*v*] will be equal to 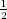. These probabilities are used instead of the genome-wide error term *∈* in equations 4, 5 and 6. For instance, Equation 4 for a specific base b in fragment *R_j_* becomes:

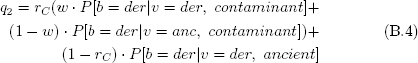

Here, *der* is the derived base and *anc* is the ancestral base. In case different post-mortem damage matrices are provided by the user for the ancient and the contaminant fragments, the events *contaminant* and *ancient* serve to denote which damage probabilities (i.e. *P_dam_*) should be used in each case.

